# Evolving Social Context Regulates Contagion Spread in Multiplex Networks

**DOI:** 10.64898/2026.06.15.732209

**Authors:** Diksha Gupta, Baltazar Espinoza

## Abstract

Contagion processes across biological, behavioral, and informational systems are shaped not only by transmission dynamics, but also by adaptive social responses emerging through human interactions. Understanding how social context regulates contagion spread is therefore critical for characterizing real-world spreading processes. Yet standard epidemic models often focus primarily on contagion states, treating social context as static or only weakly coupled to transmission. Here, we develop a multiplex-network framework that couples contagion dynamics with co-evolving social context. Unlike classical threshold contagion models, which apply thresholds directly to contagion prevalence or adoption states, our framework applies heterogeneous local and global thresholds to evolving context dynamics. The model further captures context-mediated transmission through targeted spread, in which contagion selectively propagates toward the locally most context-vulnerable susceptible individual. This contrasts with broadcast transmission, where spreading effort is distributed uniformly across susceptible neighbors. We show that coupling contagion with evolving context fundamentally reshapes spreading dynamics, producing delayed convergence and non-monotonic final prevalence. Targeted and broadcast transmission mechanisms exhibit distinct sensitivities to local and global social responses, highlighting tradeoffs in intervention strategies. We further show that co-evolving context can generate resilience by slowing propagation and delaying equilibrium, while pre-existing social resilience can substantially suppress contagion even under high transmission rates. These results suggest that contagion outcomes can vary substantially as a function of evolving social response and pre-existing social resilience.

**Significance:** Contagion outcomes are often shaped before transmission begins. Existing social environments can make populations more vulnerable or more resistant to future spread, yet this latent resilience is difficult to capture when interventions are represented only as changes to contact or transmission rates. Our results show that social context can act as a regulatory mechanism that suppresses and delays contagion spread. In particular, prior pro-social alignment can create resilience before exposure occurs, helping explain why community prevention, peer support, and reintegration programs may alter contagion outcomes even when they do not directly target the transmission process.

## Introduction

Human behavior plays a fundamental role in shaping contagion processes across diverse domains. Contagion processes are pervasive and may refer not only to biological pathogens but also to behavioral phenomena such as addiction [1, 37], emotions [3, 25, 18], or informational processes including political and marketing campaigns [24, 20, 35]. Notably, harmful behaviors such as suicide and self-harm have also been shown to exhibit contagion-like dynamics, where exposure within social networks can modulate risk shaping individual predisposition [4]. In many such settings, transmission dynamics are not solely governed by the characteristics of the contagion process, but are mediated by individual and collective behaviors embedded within social networks, whose interactions give rise to complex contagion dynamics.

Singapore’s drug-control history provides a motivating example for this modeling challenge. Although Singapore has long adopted a zero-tolerance stance toward drugs, its prevention and rehabilitation ecosystem has increasingly incorporated community-based strategies that act through social context rather than enforcement alone [27]. For example, the Singapore Anti-Narcotics Association shifted parts of its preventive strategy toward youth engagement, peer-led campaigns, online outreach, and support for ex-offenders and families, in response to concerns about younger, more affluent, and more educated drug users. This illustrates a broader feature of behavioral contagions: vulnerability is shaped not only by exposure, but also by evolving peer influence, family support, stigma, youth attitudes, and community trust.

Consequently, social interventions that target behavior and social context, rather than the contagion process directly, emerge as powerful tools for controlling spread. Examples include the seminal work by Kaplan [22] that demonstrated that expanding access to clean needles can substantially reduce HIV transmission, by altering behavioral practices without directly targeting the virus. More recently, during the COVID-19 pandemic, widespread adoption of mask-wearing [17, 32] and social distancing [36, 31], along with vaccination, similarly reduced effective transmission by modifying individual susceptibility and exposure.

Classical epidemic models primarily emphasize the role of network structure in governing contagion spread through direct contact processes. A large body of work extends classical contagion models to account for social reinforcement and behavioral adoption through *complex contagion* mechanisms. Unlike simple contagion processes, where exposure to the contagion governs transmission, complex contagions evolve as a function of an individual’s social neighborhood. Threshold-based frameworks have been widely used in literature to capture such dynamics. For instance, Granovetter [14] introduced heterogeneous adoption thresholds to explain collective behavior, while Watts [38] demonstrated how network structure and threshold distributions jointly determine cascade dynamics of contagion spread. Subsequent models have incorporated heterogeneous sensitivities and competing influences, revealing rich phenomena such as cascade fragility, critical mass effects, and non-linear adoption patterns. These approaches have been widely used to model social contagion such as opinion formation, norm diffusion, and behavioral change across social systems.

An important but comparatively underexplored dimension of contagion dynamics is the role of *transmission mechanisms* governing how influence or infection propagates across contacts. Classical epidemic and network-based models typically assume homogeneous transmission processes, where an infected individual interacts uniformly with all neighbors through fixed per-contact probabilities [30]. In contrast, real-world spreading processes often exhibit heterogeneity in how influence is allocated, ranging from broadcast-like interactions that distribute attention or exposure across many contacts, to targeted interactions that focus influence on specific individuals. Although prior work has examined heterogeneity in transmission rates and contact structures, the underlying *mechanism of transmission* itself is usually treated as implicit and fixed. As a result, existing models provide limited insight into how different modes of influence allocation shape contagion dynamics, particularly in settings where social context co-evolves with contagion dynamics.

Recent work investigates the interplay between social behavior and contagion dynamics through coupled and multilayer models. Studies of vaccination behavior and epidemic spread show that behavioral responses can significantly alter outbreak size and timing by introducing feedback between perceived risk and transmission dynamics [34, 11]. Related work on awareness diffusion and adaptive behavior demonstrates that information spread can suppress or reshape epidemics, leading to phenomena such as multiple waves, bistability, and network fragmentation [15, 10, 12, 13]. More broadly, multilayer and multiplex network models capture interactions between distinct spreading processes across different channels of interaction, highlighting how cross-layer coupling influences diffusion outcomes [21, 6].

While these models incorporate feedback between social and contagion processes, they often treat social influence as a fixed variable that affects the contagion dynamic, rather than modeling the coevolution of the social behavior along with the contagion. A deeper understanding of these mechanisms is crucial for designing targeted and effective intervention strategies. Moreover, these studies majorly modeled behaviors based on homogeneous thresholding approaches, where sensitivities of the population are assumed to be well-aligned with each other. These assumptions can produce population-wide cascades once a critical mass is reached, but may underrepresent variability in individual responses [16, 32].

Our framework introduces a latent social-context layer with heterogeneous thresholding mechanisms that operate based on the state of the social-context instead of the classic contagion state based mechanisms. Moreover, we consider non-uniform thresholding mechanisms at individual level enabling asynchronous activation and multi-scale feedback that fundamentally alters contagion dynamics.

### Our Contributions

In this work, we develop a social-context coupled contagion framework that captures the coevolution of multi-scale social dynamics and contagion processes. The model consists of two interacting layers over the same population: a context layer encoding evolving individual beliefs, and a contagion layer governed by susceptible-infected (SI) dynamics. The context layer captures the interplay between pro-social and contagion-aligned influences through a continuous context score that evolves via local peer-pressure thresholding and global social-conformity thresholding, inducing a dynamic social field that both shapes and responds to contagion spread. The contagion layer retains the simplicity of the SI model while modulating transmission probabilities as a function of individual context, thereby introducing feedback between social states and observed spread; we further incorporate both broadcast and targeted transmission mechanisms. Despite this simple SI backbone, the coupled system exhibits rich emergent behavior, enabling us to isolate and systematically study the impact of behavioral factors on contagion dynamics. Our formulation generalizes a broad class of existing models, recovering classical epidemic models in the absence of social modulation and threshold-based behavioral models when contagion dynamics are suppressed. By explicitly modeling the coevolution of social context and contagion, the framework provides a unified lens for studying socially mediated contagions, including addiction-like processes where peer influence, community norms, and reintegration support jointly shape vulnerability. We derive the following insights based on our framework:

- **Contagion transmission characteristics govern contagion spread, exposing tradeoffs between global and local response**. We focus on understanding the difference in contagion spread dynamics as a function of the underlying contagion transmission mechanism i.e., whether the contagion spreads uniformly to all its neighbors via broadcast or it spreads through focused transmission via targeting a single neighbor. We observe that targeted contagions with high transmission rates are more sensitive to local interventions in the model, whereas global interventions are more effective in curbing the spread of low rate infections.
- **Pro-social leaning context resists contagion, delaying convergence to equilibrium**. We model social context as a layer that jointly evolves with the contagion dynamics as a function of both the ongoing infection spread and the existing population social profiles within the network. This co-evolution gives rise to social resistance arising from pro-social sentiment that creates transmission barriers to contagion propagation, making it increasingly difficult for the contagion process to infect individuals and thereby delaying the convergence of the system to equilibrium.
- **Pre-existing pro-sociality coupled with heterogeneous sensitivities, regulates contagion spread via dispersed adoption**. We show that due to variable individual sensitivity thresholds, social context evolves slowly and non-uniformly over the population resulting in rich contagion dynamics. Moreover, existing pro-social sentiment can effectively dampen contagion spread even at high transmission probability, pointing to the role of long-term social campaigns against infections. In policy terms, this suggests that prevention campaigns and peer-support structures may matter not only during active spread, but also by shaping the initial social-context distribution before exposure occurs.

### Methodology

We model the contagion process using a social-context coupled multiplex network, illustrated in Figure 1. Our system consists of two layers defined over the same set of individuals *V*, which capture two types of interactions among individuals:

**Figure 1:**
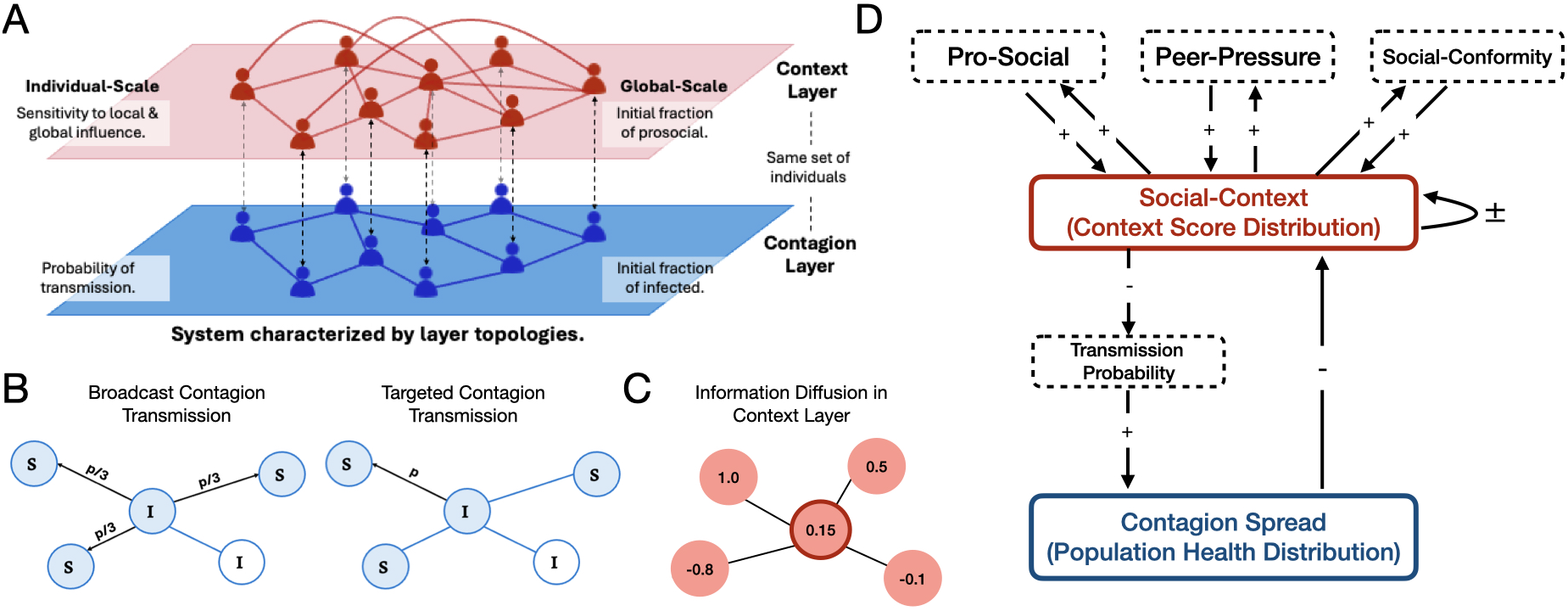
Schematic of the mechanisms incorporated in our framework. (A) Multi-scale framework for coupled social-context and contagion dynamics. The context layer (top) captures behavioral influences via local peer effects, global conformity, and initial prosocial fraction, while the contagion layer (bottom) models infection spread with transmission probability and initial infections. The layers are coupled at the individual level, enabling feedback between social context and contagion dynamics. (B) The proposed framework consists of a contagion layer, in which an infected individual spreads infection through either *broadcast contagion transmission*, where the infection probability is distributed uniformly across all neighbors, or *targeted contagion transmission*, where infection is directed to a selected neighbor with the full transmission probability. (C) Additionally, the social-context is maintained as a score for each individual in the range [−1, 1] that spreads in the context layer through diffusion as average of neighboring scores. (D) The flow chart illustrates overall interaction among the modeled components from (A), (B) and (C), governing the dynamics of the context and contagion layers.

- The **Context layer**, capturing the role of weak-ties such as the social media interactions and acquaintances, through which social influence and contextual information diffuse. Each individual *i* maintains a continuous *context score*, 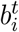 in the interval [−1, 1], representing its attitude toward the contagion. The context score evolves as a function of its neighboring context score through diffusion and also via thresholding mechanisms described later in this section.
- The **Contagion layer**, capturing the role of close relationships of an individual such as family, close friends, trusted peers, through which the contagion spreads. Each individual *i* has a state 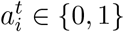, where 0 corresponds with the susceptible state, and 1 with the infected state. The contagion spread rate is modulated by the context score, which in turn evolves as individuals get infected by the contagion.

The contagion and social-context co-evolve through a feedback mechanism, where contagion exposure polarizes individual context, while evolving context dynamically regulates future susceptibility to contagion. Concretely, let *N* (*i*) denote the neighborhood of individual *i* and *p*^*t*+1^(*j, i*) denote the probability of contagion transmission from individual *j* to *i*, then we define the effective probability of a susceptible individual *i* being infected at time *t* + 1 as:

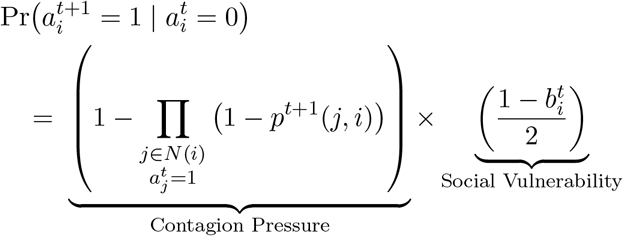

Thus, unlike classical contagion models with static susceptibility, individual vulnerability evolves dynamically as a function of the social context in our framework. Moreover, we propose a novel sentiment based social thresholding, inspired from [14, 38], where behavioral polarization is driven by perceived social sentiment rather than the infection prevalence itself. This captures the role of prevailing social sentiment to the contagion over the actual contagion state of the system as the driving factor for individual response to contagion. We model two forms of individual-level social thresholding: *Peer Pressure* denoted by *τ*_*p*_(*i*), capturing alignment with the immediate neighborhood; and *Social Conformity* denoted by *τ*_*s*_(*i*), capturing alignment with the prevailing global social sentiment. Note that we restrict *τ*_*s*_(*i*), *τ*_*p*_(*i*) ∈ [0, 1] so that polarization toward the pro-social state occurs only when the corresponding sentiment is sufficiently positive.

At each time step, the context layer is updated in three stages. First, infected individuals are polarized toward the contagion-aligned state, *b*_*i*_ = −1. Second, susceptible individuals update their context score by averaging over their context-layer neighborhood. Third, local peer-pressure and global social-conformity thresholds polarize individuals toward the pro-social state, *b*_*i*_ = +1, when neighborhood-level or population-level sentiment exceeds individual thresholds. Thus, social context both responds to contagion exposure and regulates future susceptibility. Further details are provided in the Supplementary Information. Note that in all simulations, the context update is applied before the contagion update within each time step.

We consider two models of contagion transmission. Let 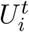 denote the set of susceptible neighbors of individual *i* in the contagion layer at time *t*, and let *p* denote the baseline transmission probability.

- **Broadcast transmission**. In this model, an infected node distributes its transmission probability uniformly across all susceptible neighbors. Thus, the probability that node *i* transmits the contagion to a susceptible neighbor *k* is

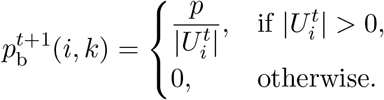
- **Targeted transmission**. In this model, an infected node preferentially targets the most context-vulnerable susceptible neighbor, with ties broken uniformly at random. The corresponding transmission probability is

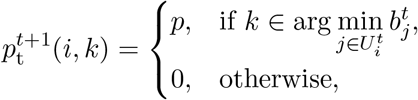

where 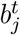 is the context score of susceptible individual *j* at time *t*. Lower context scores therefore indicate greater vulnerability to the contagion.

Although the main analysis uses an SI backbone to isolate the regulatory role of evolving social context, the framework is not restricted to irreversible contagion. In the Supplementary Information, we also consider an SIR extension under broadcast transmission, where infected individuals recover after a fixed recovery duration and then remain immune. To keep the recovery mechanism consistent with the social-context layer, recovered individuals are treated as pro-social, with their context score set to +1 after recovery. Further details are provided in the Supplementary Information.

### Experimental Setup

Unless otherwise stated, simulations use a Barabási–Albert (BA) network with 30, 000 individuals, an initial infected fraction of 0.003, an initial pro-social fraction of 0.01, and results averaged over 100 random simulation runs; full parameter details and robustness experiments are provided in the Supplementary Information.

## Role of Contagion Transmission mechanism in co-evolving Social Context and Contagion Dynamics

In this section, we begin by contrasting contagion behavior of the susceptible-infected (SI) model under various context-coupled SI model scenarios (Fig. 2). In the absence of social-context, corresponding to the *No Context Scenario*, contagion spreads monotonically with transmission probability *p*, and convergence time decreases smoothly, reflecting standard SI dynamics. In contrast, introducing the context layer fundamentally reshapes both the final contagion fraction and the convergence behavior of spreading processes.

**Figure 2:**
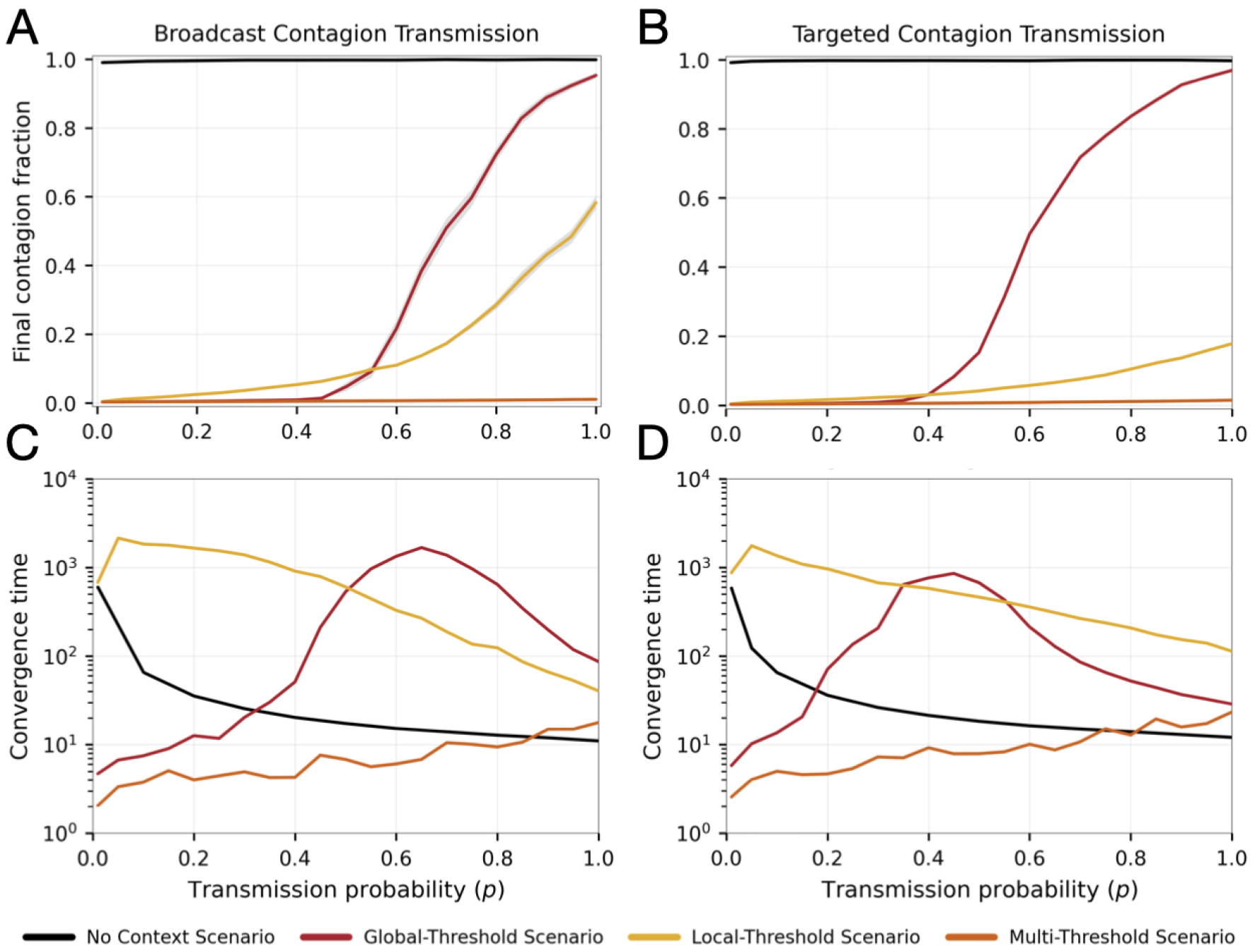
Heterogeneous multi-scale social-context results in a variety of emergent contagion dynamics, even under a simple SI model. We compare (A-B) final contagion fractions and (C-D) Convergence time under broadcast and targeted transmission for the baseline SI model and four context-threshold variants. The *No Context Scenario* refers to the simple SI model without any social-context. The *No Threshold Scenario* closely mimics the SI model but with delayed convergence, due to resistance emerging from pro-social context despite the absence of any polarizing thresholds. The *Global-Threshold Scenario* modulates context using system-wide social conformity only, the *Local-Threshold scenario* uses peer-pressure from immediate neighborhood context only, and the *Multi-Threshold Scenario* incorporates both local and global thresholding.

First, despite the absence of polarization in the No Threshold Scenario, we observe an increase in convergence time relative to the No Context Scenario, widening as the transmission probability increases, while the final contagion fraction remains unchanged. Next, the Global-Threshold Scenario exhibits pronounced nonlinearities, including abrupt increases in contagion prevalence once a critical threshold is crossed. This transition is accompanied by a pronounced peak in convergence time, indicating critical slowing down near the tipping point pointing to increased systemic resistance to the contagion induced by the social context.

In contrast, the Local-Threshold Scenario produces gradual and limited contagion spread, with consistently high convergence times driven by fragmented transmission under heterogeneous local resistance with lower contagion prevalence in targeted contagion transmission. This suggests that localized responses to rapidly spreading targeted contagion can sharply reduce prevalence while greatly slowing down the spread. Finally, the Multi-Threshold Scenario further suppresses contagion, stabilizing rapidly to a low-contagion regime. These behaviors persist across both broadcast and targeted transmission mechanisms.

Importantly, contagion outcomes are no longer monotonic in transmission rate: moderate values of *p* can produce slower spread than low-transmission regimes and higher prevalence than high-transmission regimes, depending on the social response to the contagion. Together, these results demonstrate that coupling a simple SI process with heterogeneous multi-scale thresholding induces rich nonlinear phenomena, including critical transitions, and suppression regimes. More broadly, the results highlight the role of latent social-context as a regulatory layer governing both the speed and extent of contagion spread.

This distinction has a natural analogue in drug-prevention settings. Broad anti-drug messaging, online engagement, and school-wide campaigns resemble broadcast-facing interventions, whereas programs for at-risk youths or peer-group disruption operate more locally. Singapore’s community-based prevention efforts illustrate this combination: SANA expanded digital and youth engagement while also piloting targeted decision-making programs for at-risk students, suggesting that intervention scale may need to be matched to the way exposure is allocated through the social network.

## Temporal Regulation of Contagion Dynamics

We next examine the temporal dynamics underlying the emergent contagion regimes introduced by heterogeneous social-thresholding (Fig. 3). While Fig. 2 characterizes equilibrium outcomes and convergence behavior, the temporal trajectories reveal the mechanisms through which evolving social-context regulates contagion spread.

**Figure 3:**
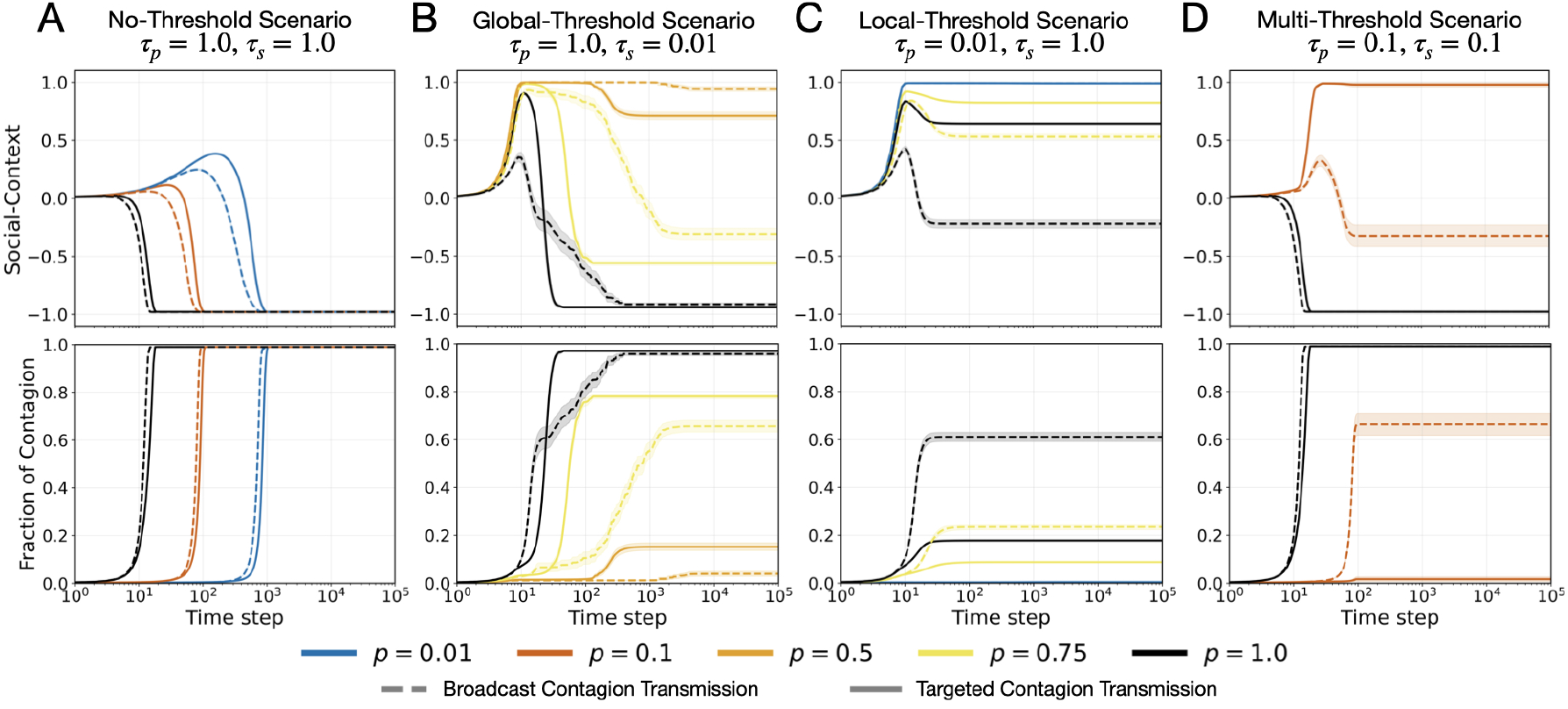
SI Temporal dynamics of the context and contagion layers under heterogeneous social-thresholding regimes. We illustrate the evolution of the *social-context*, computed as the average context score across all individuals in the system, and the *fraction of contagion*, computed as the fraction of infected individuals at each time step. We compare the temporal trajectories across a range of transmission probabilities, *p*, and contrast broadcast and targeted contagion transmission mechanisms under different social-thresholding regimes. The results demonstrate that heterogeneous social-thresholding can induce emergent social resistance, suppress contagion spread, and significantly delay convergence to equilibrium.

In the No-Threshold Scenario, the coupled framework closely mimics baseline SI dynamics, but convergence becomes delayed due to resistance emerging from pro-social context despite the absence of explicit polarization thresholds. In contrast, the Global-Threshold Scenario exhibits a characteristic two-stage dynamic. Contagion must first accumulate sufficiently to shift the global social-context before transmission becomes effective, resulting in an initial suppression phase followed by a rapid cascade once enough conformity thresholds are crossed. This delayed activation mechanism explains the critical slowing down and abrupt phase transitions observed in Fig. 2.

The Local-Threshold Scenario instead produces fragmented and heterogeneous contagion trajectories, where local peer-pressure creates persistent resistance pockets that suppress large-scale synchronization. Consequently, contagion spread remains gradual and limited even at moderate or high transmission probabilities. The Multi-Threshold Scenario amplifies these suppression effects further, rapidly stabilizing the system in regimes dominated by pro-social context, thereby preventing widespread contagion propagation.

Importantly, unlike much of the threshold-based contagion literature where thresholds are concentrated near a critical value, our framework samples thresholds uniformly from (*τ*_*x*_, 1), inducing substantial heterogeneity in both local and global social responses. This broader threshold distribution prevents synchronized activation and generates staggered adoption dynamics, fragmented spreading, and sharp tipping behavior across transmission regimes.

Together, these temporal dynamics demonstrate that system-level complexity emerges not from the contagion process alone, but from the interaction between contagion dynamics and heterogeneous, evolving social-context. Even under a simple SI backbone, the coupled framework produces delayed cascades, suppression regimes, and nonlinear temporal transitions driven by multi-scale behavioral feedback. This delayed-convergence behavior is consistent with settings where behavioral or social norms reduce effective susceptibility without eliminating the underlying contagion source. Examples include needle-exchange participation, masking, and social distancing in public-health settings; in addiction-related settings, peer-support and reintegration programs can similarly create social resistance barriers that slow relapse or behavioral spread.

## Response Implications of Pre-existing Social profiles

We next investigate how pre-existing social conditions regulate contagion dynamics across heterogeneous thresholding regimes (Fig. 4). Specifically, we analyze the final contagion fraction and convergence time as functions of the transmission probability *p* and the initial fraction of pro-social individuals in the population. Across both local and global thresholding regimes, increasing the initial pro-social population progressively shifts the system from a high-contagion regime toward suppression, revealing sharply defined contagion boundaries separating these two dynamical phases.

**Figure 4:**
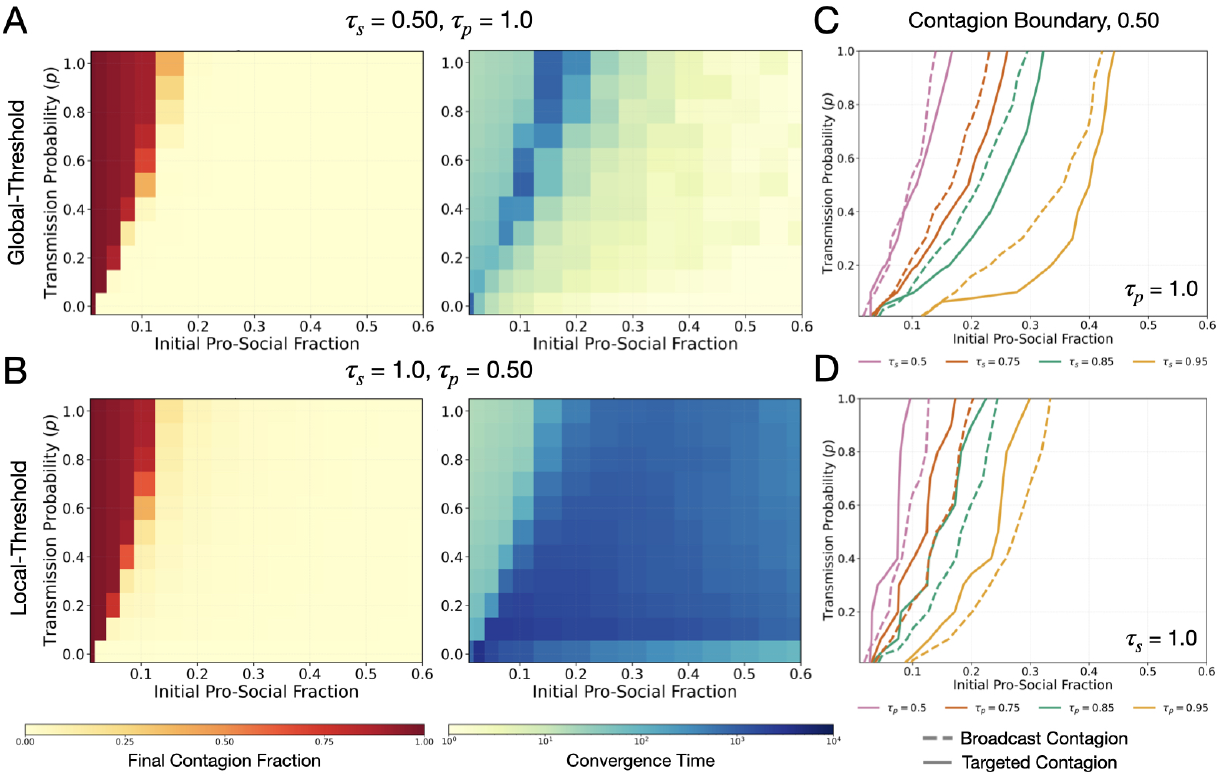
Impact of pre-existing pro-social sentiment on contagion dynamics. We analyze the *final contagion fraction* and the *convergence time* required to reach equilibrium across varying transmission probabilities, *p*. (A-D) We compare global and local social-thresholding regimes under similar sensitivity values, illustrating how different individual social responses alter contagion dynamics, despite comparable initial pro-social sentiment. (E-F) We illustrate the contours corresponding to a final contagion fraction of 0.5 as a function of the initial pro-social fraction and transmission probability, *p*. Dashed curves correspond to broadcast contagion transmission, whereas solid curves correspond to targeted transmission. The results demonstrate that increasing social-threshold sensitivity shifts the critical contagion boundary toward needing larger pro-social sentiment irrespective of contagion characteristics.

The nature of this transition, however, depends strongly on the scale of social influence. Under the Local-Threshold Scenario, contagion boundaries evolve gradually with transmission probability, reflecting the heterogeneous and decentralized influence of peer-pressure. Increasing the initial pro-social population incrementally suppresses contagion by generating localized resistance pockets that fragment spreading pathways. Consequently, the transition between contagion and suppression regimes remains diffuse, with convergence times remaining elevated across broad regions of parameter space.

In contrast, the Global-Threshold Scenario produces substantially sharper and more coherent phase transitions. Small increases in the initial pro-social population can abruptly suppress contagion across a wide range of transmission probabilities, indicating strong collective sensitivity to global conformity dynamics. Temporal convergence patterns further reveal the presence of critical slowing down near the phase boundary, suggesting that contagion must first accumulate sufficiently to shift the global social-context before widespread activation becomes possible.

The resulting contagion boundaries, illustrated in Fig. 4(E-F), demonstrate that increasing social-threshold sensitivity systematically shifts the critical contagion frontier toward requiring larger transmission pressures and lower pro-social alignment for sustained spreading. Moreover, the distinction between broadcast and targeted transmission becomes increasingly pronounced near these tipping regions, with targeted contagion typically producing sharper transitions and narrower activation boundaries once local resistance barriers are breached.

Together, these results demonstrate that pre-existing social conditions act as a latent regulatory layer governing contagion outcomes. Even under a simple SI backbone, heterogeneous thresholding transforms contagion spread into a phase-dependent process characterized by tipping behavior, delayed activation, and suppression regimes. More broadly, the findings suggest that interventions promoting pro-social alignment can induce system-wide reductions in contagion by shifting populations across critical dynamical boundaries.

We next examine how heterogeneous social-response profiles jointly regulate contagion dynamics under the Multi-Threshold Scenario (Fig. 5). Unlike the isolated local or global thresholding regimes, the coupled framework integrates both peer-pressure and social-conformity mechanisms, producing rich nonlinear contagion landscapes governed by competing local and global feedback processes.

**Figure 5:**
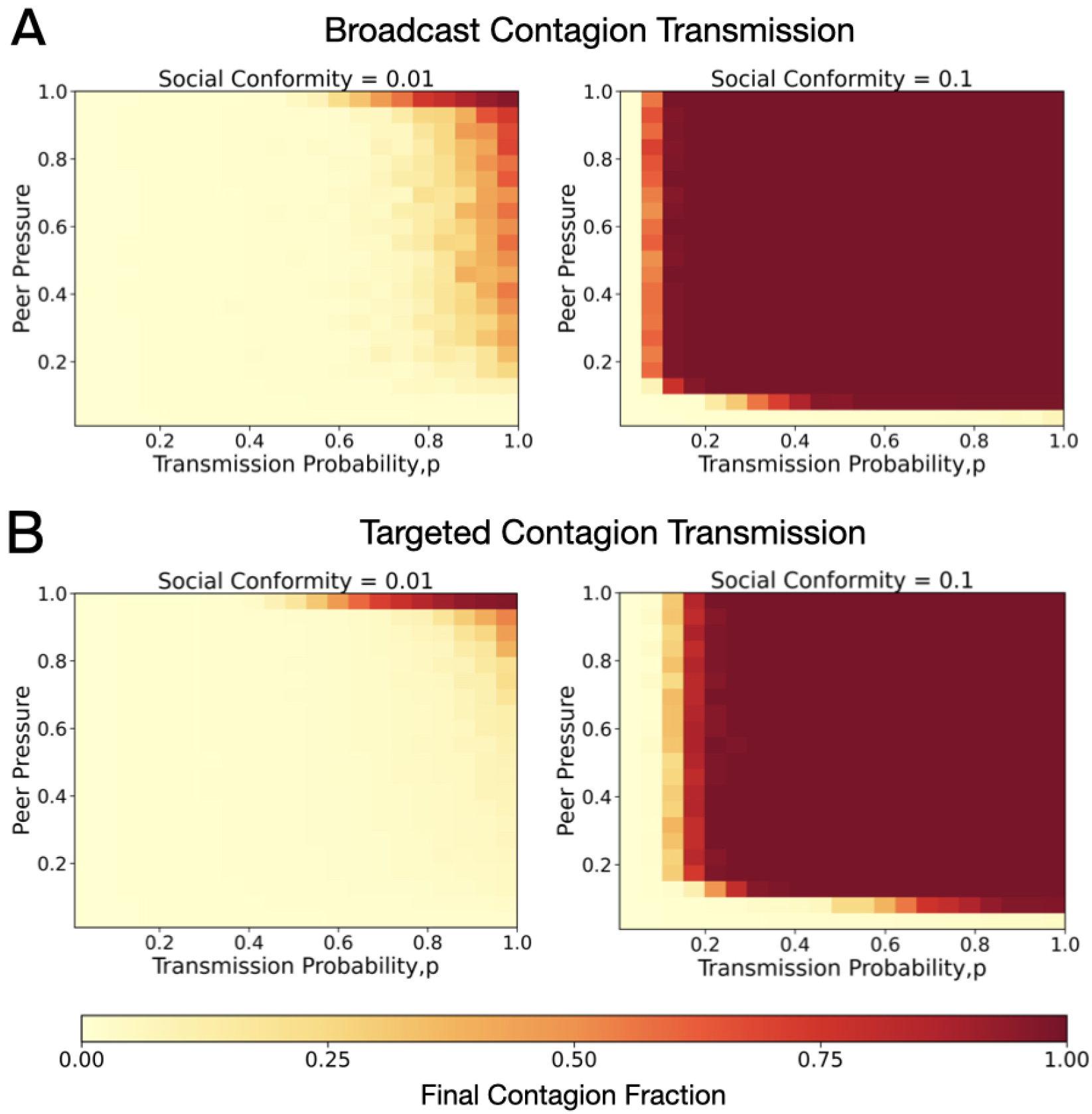
Impact of social-response profiles on the emergent contagion in Multi-Threshold Scenario. We illustrate the impact of varying social-thresholding profiles on the final fraction of infected population at equilibrium as a function of contagion characteristics: the contagion transmission probability and transmission mechanism.

Increasing local peer-pressure thresholds *τ*_*p*_ progressively suppresses contagion by strengthening localized resistance to spreading. For low *τ*_*p*_, contagion propagates readily across the network, whereas larger threshold values increasingly fragment transmission pathways, shifting the critical transition toward higher transmission probabilities. In contrast, varying global social-conformity thresholds *τ*_*s*_ produces a complementary collective effect. Lower *τ*_*s*_ values facilitate rapid global alignment, enabling large-scale contagion cascades once conformity thresholds are crossed, while higher *τ*_*s*_ delays or prevents coordinated activation, confining contagion to localized regions. Moreover, due to information diffusion in the context layer, the social-context creates higher prosocial-population fraction faster than targeted contagion transmission spread, limiting the spread of contagion in comparison to high broadcast contagion where the contagion and context spread at comparable strength.

Importantly, the interaction between *τ*_*p*_ and *τ*_*s*_ generates strongly non-separable contagion dynamics that cannot be explained by either mechanism independently. Small changes in threshold values produce abrupt transitions between suppressed and high-contagion regimes, revealing tipping behavior emerging from multi-scale social feedback. These regime boundaries further differ across broadcast and targeted transmission mechanisms, demonstrating that contagion characteristics and social-response heterogeneity jointly determine spreading outcomes.

From an intervention perspective, these results suggest that distinct scales of social influence produce qualitatively different mitigation effects. Increasing local resistance through peer-based interventions can suppress contagion even at moderate transmission probabilities, whereas modifying global perception or social norms can inhibit large-scale cascades by preventing collective alignment. The presence of sharp phase boundaries further implies that relatively small but well-targeted interventions may shift the system across critical thresholds, leading to disproportionate reductions in contagion prevalence.

This result highlights the importance of interventions that act before exposure becomes widespread. Youth ambassador programs, peer-led anti-drug campaigns, and community trust-building efforts can be interpreted as attempts to reshape the initial social-context distribution rather than only responding after contagion has emerged. In the Singapore case, SANA’s youth engagement strategy explicitly empowered students to organize peer-facing drug education campaigns, while the report emphasizes that community approaches require sustained trust, capability building, and social capital rather than one-off interventions.

## Conclusion

Classical epidemic models suggest that increasing transmission probability monotonically increases contagion spread. Our results demonstrate that this intuition fundamentally breaks down when contagion is coupled with heterogeneous, multi-scale social-context. Even under a simple susceptible-infected (SI) process, contagion outcomes become strongly non-monotonic, with moderate transmission regimes exhibiting delayed spread or even lower equilibrium contagion than more infectious settings. These dynamics emerge from feedback between contagion and evolving social-context, where transmission must first overcome locally and globally mediated resistance before widespread cascades become possible.

Across the coupled framework, we identify three distinct dynamical regimes induced by multi-scale thresholding: (i) a suppression regime, where strong pro-social alignment or localized resistance prevents contagion propagation; (ii) a transition regime characterized by delayed activation, fragmented spreading, and critical slowing down; and (iii) a saturation regime in which contagion rapidly propagates once social thresholds are exceeded. These regimes are separated by sharp phase boundaries, revealing strong sensitivity to both initial social conditions and the balance between local peer-pressure and global social-conformity dynamics.

From an intervention perspective, our findings suggest that modifying transmission alone may not produce predictable outcomes when social-context co-evolves with contagion. Addressing transmission without reinforcing pro-social norms can delay contagion while leaving the system vulnerable to abrupt resurgence once critical thresholds are crossed. Importantly, interventions acting at different social scales produce qualitatively distinct effects: local interventions increase resistance and fragment spreading process, whereas global interventions reshape collective alignment and can suppress large-scale cascades. Their interaction is inherently nonlinear, implying that combined interventions may generate synergistic, delayed, or counterintuitive outcomes.

More broadly, our results demonstrate that social-context functions as a latent regulatory layer fundamentally altering contagion dynamics. System-level complexity emerges not from the contagion process alone, but from its interaction with heterogeneous and evolving social environments. This perspective provides a unifying framework for understanding spreading phenomena across biological, behavioral, and informational systems, while underscoring the importance of incorporating adaptive social dynamics into contagion modeling and intervention design. Moreover, our findings are robust to relaxing the irreversible SI assumption. In the Supplementary Information, we consider an SIR extension with finite recovery durations of 15 and 180 days. The results show that pro-social context substantially suppresses short-duration contagions, while longer recovery durations reveal tradeoffs between local and global interventions. This suggests that the proposed mechanism is not an artifact of the SI backbone; evolving social context remains a regulatory layer even when individuals recover and acquire immunity.

## Acknowledgment

We thank members of BI for a number of useful comments and discussions. This material is based upon work supported by the National Science Foundation under Grant No. CNS-2327710 and the UVA Contagion Science program.

## SUPPLEMENTARY MATERIAL

In this section, we detail the existing literature on opinion spread in networks, followed by details of our multi-layer contagion models and additional experimental details & findings.

### A Related Work

#### Complex contagions and threshold models

A large body of work models social behaviors—such as opinion formation, norm adoption, and protective actions—as *complex contagions*, whose spread requires reinforcement from multiple exposures rather than single contacts. Centola and Macy [7] formalized this distinction and demonstrated that network structure can fundamentally alter diffusion outcomes, with long ties sometimes inhibiting rather than facilitating spread. Threshold models provide a canonical abstraction of complex contagions. Granovetter [14] introduced individual adoption thresholds to explain collective behavior, while Watts [38] extended this framework to random networks and identified conditions under which small initial perturbations can trigger global cascades. Subsequent work has examined heterogeneous thresholds, degree effects, and cascade fragility, establishing threshold dynamics as a foundational model of social diffusion.

Several extensions of threshold models incorporate additional mechanisms such as memory, persuasion, reversibility, and competing influences. Dodds and Watts [8, 9] proposed a generalized contagion framework blending independent-cascade and threshold mechanisms, identifying distinct dynamical regimes including epidemic thresholds and critical mass effects. Other studies have introduced reversible adoption, bi-threshold dynamics, and persuasion effects to capture asymmetries between adoption and abandonment and the role of highly influential agents [33, 26, 19]. These models have been applied across domains including smoking behavior, coordination games, and innovation diffusion, highlighting the richness of threshold-based social dynamics.

#### Multilayer contagion models

Multiplex and multilayer network formulations provide a natural framework for modeling interacting contagions. Prior work shows that diffusion across multiple layers can accelerate or suppress spreading depending on cross-layer coupling strength and structural heterogeneity [6, 21]. While these models capture structural interactions between layers, they often assume fixed transmission rules within each layer and do not explicitly model the evolution of latent social variables that regulate contagion acceptance.

Most closely related to our approach are recent network-based models that allow social behavior and epidemics to coevolve concurrently. In particular, Qiu et al. [32] demonstrate that coupling mask-wearing behavior with disease dynamics can generate robust critical transitions driven by behavioral feedback. Our framework generalizes this line of work in three key ways. First, individuals maintain a continuous latent social-context state alongside a discrete contagion state, allowing belief, vulnerability, and behavior to coevolve. Second, social context evolves through explicit local averaging and nonlinear polarization mechanisms capturing peer pressure and conformity, rather than purely probabilistic adoption. Third, contagion transmission is endogenously regulated by social context and can operate under distinct mechanisms, including targeted transmission based on social alignment. Together, these features unify threshold-based social contagions and epidemic spreading within a single co-evolving framework, enabling the analysis of socially mediated interventions and feedback-driven transitions that are not captured by existing models.

#### Coupled social and contagion dynamics

A related line of work investigates the concurrent spread of social behaviors and biological diseases on networks. Early models focused on vaccination behavior coupled with epidemic dynamics, showing that behavioral responses can substantially alter outbreak size and timing [11]. Subsequent studies examined how opinions about vaccination or preventive behaviors cluster socially and feed back into disease spread [34, 5]. These models demonstrate that coupling social and biological processes can generate nonlinear outcomes, including critical transitions, bistability, and hysteresis effects. Other approaches emphasize awareness, fear, and behavioral adaptation during epidemics.

Models of awareness diffusion show that information spread can dampen infection rates and induce oscillations or multiple epidemic waves [12, 13]. Related work investigates fear propagation, behavioral withdrawal, and adaptive network rewiring as responses to perceived risk, demonstrating that behavioral adaptation can fragment contact networks or reshape epidemic thresholds [10, 15]. These studies highlight the importance of feedback between perceived risk and disease dynamics, but often treat social influence as a global signal rather than an endogenously evolving social process.

#### Targeted transmission in behavioral contagions

Prior work on information diffusion has extensively studied *targeted influence*, primarily from the perspective of selecting individuals or groups to maximize spread. The influence maximization framework [23] formalizes the problem of identifying a set of seed nodes that can trigger large cascades, and subsequent work has extended this paradigm to target specific subpopulations or optimize influence under competing campaigns [28, 39]. While these approaches capture important aspects of targeted spreading, they primarily focus on *who* should be influenced or *which populations* should be reached.

A related line of work incorporates selective propagation through content relevance, topical affinity, or external sources of influence, distinguishing peer-to-peer diffusion from broadcast mechanisms such as mass media [29, 2]. However, these models generally leave implicit the individual-level *mechanism of transmission*: how an infected individual allocates influence across its contacts during propagation. As a result, existing contagion models typically do not distinguish between broadcast-like processes that distribute influence broadly across neighbors and targeted processes that concentrate influence on specific contacts. Our work addresses this gap by explicitly modeling targeted transmission as a contagion mechanism, in which exposure is directed toward the most context-vulnerable susceptible neighbor rather than spread uniformly across all neighbors.

### B Motivating Historical Scenario: Singapore Drug-Control Policy

Singapore provides a useful motivating example for studying behavioral contagion under evolving social context. Its drug-control regime has historically combined strong legal deterrence with prevention, rehabilitation, and reintegration efforts. While the country is widely known for its zero-tolerance stance toward drugs, its policy history also illustrates that drug use and relapse are not governed solely by individual exposure or enforcement pressure. Rather, they are shaped by social attitudes, peer influence, family support, community acceptance, and the broader institutional environment in which individuals are embedded.

This historical setting motivates our social-context coupled contagion framework in three ways. First, Singapore’s experience highlights the importance of distinguishing the *state of contagion* from the *social context* surrounding that contagion. Drug use may be treated as the contagion-like behavior, but the vulnerability of an individual to initiation or relapse depends strongly on attitudes toward drugs, stigma, perceived social acceptance, and exposure to supportive or criminogenic peer environments. This motivates our use of a continuous context score 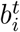, which captures an individual’s evolving attitude toward the contagion and regulates susceptibility independently of infection prevalence alone.

Second, the Singapore case illustrates the multi-scale nature of social influence. Local peer environments matter: negative peer groups can increase relapse risk, while positive peer networks, peer leaders, family support, and community spaces can reinforce recovery. At the same time, system-wide sentiment also matters: more liberal attitudes toward drugs, online narratives, and changing youth perceptions can alter the background social climate in which prevention and relapse occur. This motivates our separation between *peer pressure*, modeled through local neighborhood sentiment, and *social conformity*, modeled through global social sentiment. In this sense, behavioral polarization in our framework is driven not merely by the number of infected individuals, but by the perceived social meaning of the contagion.

Third, Singapore’s community-based prevention and rehabilitation efforts motivate our distinction between broadcast and targeted transmission. Traditional public education campaigns resemble broadcast processes, where messages or influences are distributed broadly across a population. By contrast, targeted outreach to at-risk youths, peer-led support groups, and reintegration programs directed toward individuals in vulnerable transition periods resemble targeted processes, where influence is concentrated on specific individuals whose social context makes them more susceptible to either relapse or recovery. Our targeted transmission model abstracts this mechanism by allowing an infected individual to concentrate transmission on the most context-vulnerable susceptible neighbor, rather than distributing influence uniformly across all contacts.

The Singapore case also motivates the feedback structure of the model. Contagion exposure can reshape individual context: relapse, recovery, stigma, or successful reintegration may change how individuals perceive the behavior and how they are perceived by others. Conversely, the evolving context regulates future vulnerability by either amplifying susceptibility through negative peer influence and permissive attitudes, or suppressing contagion through pro-social sentiment, supportive networks, and community reintegration. Thus, the framework is designed to capture a central lesson from the Singapore setting: behavioral contagions unfold not only through contact networks, but through the co-evolution of exposure, social sentiment, and targeted social response.

### C Modeling framework

We model the spread of a contagion over social-context-coupled multiplex network. Our system consists of two layers defined over the same set of individuals *V*, which capture the influences of two types of connections among individuals:

- The **Context layer**, capturing the role of weak-ties such as the social media interactions and acquaintances, through which individuals encounter and refine context. Each individual *i* maintains a continuous *context score*, 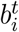 in the interval [−1, 1], representing its attitude toward the contagion, and this score evolves as a function of both the global social context and its local neighborhood. Let *G*_*s*_ = (*V, E*_*s*_) denote the network graph for the Context Layer, where *E*_*s*_ is the set of edges capturing the weak-ties among the individuals.
- The **Contagion layer**, capturing the role of close relationships of an individual such as family, close friends, trusted peers, through which the contagion spreads. Each individual *i* has a state 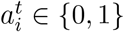 with 1 corresponding with the infected state, and 0 corresponding with the susceptible state. The infection attempts to spread along strong ties, which gets modulated by the context scores evolving over the Context Layer. Let *G*_*c*_ = (*V, E*_*c*_) denote the network graph for the contagion layer, where *E*_*c*_ is the set of edges capturing the strong-ties among the individuals.

Using the above definition of framework layers, we can now concisely define our network dynamical model, denoted by 𝒮 = (ℒ, ℱ) such that:

- ℒ = (*G*_*s*_, *G*_*c*_) is the two layer network defining the social interactions of the individuals at different scales described above, and
- ℱ denotes the update functions governing context evolution and contagion spread.

At any given time *t*, an individual *i* ∈ *V* is associated with two values: the *contagion state*, 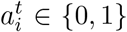; and the *context score*, 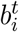 in the continuous interval [−1, 1]. Next, we define the update rules for the two layers.

#### Update rules for the Context Layer

We extend threshold-based dynamics from [14] and to capture the evolving social dynamics in the Context Layer. At each time step *t* + 1, the context score of individual *i* ∈ *V* is polarized to mimic the infected state in the contagion layer as:

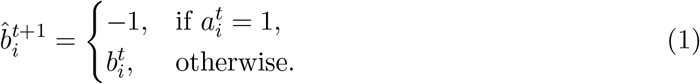

Next, we compute the mean neighborhood context score for each individual, while ensuring that infected individuals remain fixed at −1. Formally, at time step *t* + 1 for all individual *i* ∈ *V* :

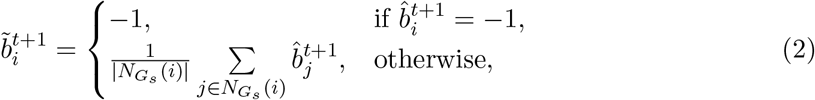

where 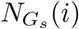 is the neighborhood of individual *i* in social graph of the Context Layer, *G*_*s*_. We assume each individual to be its own neighbor. Moreover, we consider two factors influencing individual decisions in polarizing their context. We equate this to an individual’s sensitivity to pro-social influence from their surroundings. We model two forms of social influences:

- **Peer-Pressure:** We capture the role of pro-social peer-pressure in polarizing an individual. For an individual *i*, let *τ*_*p*_(*i*) ∈ [0, 1] denote its peer-pressure thresholds such that:

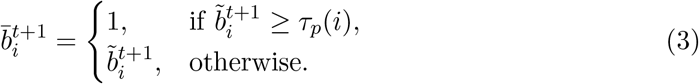
- **Global Conformity:** We capture need for pro-social alignment with the prevailing system-wide sentiment. Let 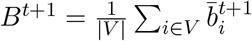 be the system sentiment towards the contagion. For an individual *i*, let *τ*_*s*_(*i*) denote the global-conformity threshold such that:

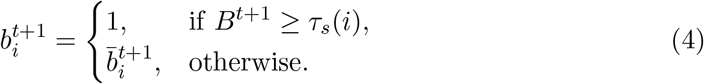

Thus, each context update consists of three stages: (i) contagion-induced polarization, (ii) local averaging over the context network, and (iii) threshold-based polarization driven by local peer pressure and global conformity. Note that we restrict *τ*_*s*_(*i*), *τ*_*p*_(*i*) ∈ [0, 1] so that polarization toward the pro-social state occurs only when local or global sentiment is sufficiently positive.

#### Update rules for the Contagion Layer

We model the spread of a contagion *I* using the SI model on *G*_*C*_. Let *p* denote the baseline transmission probability of the contagion. At any time *t*, an individual *i* ∈ *V* has state 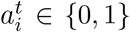, where 1 corresponds to the infected state, while the state 0 corresponds with the susceptible state. Once acquired, contagion is permanent i.e., once 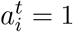, the state never reverts. Each infected individual uses one of two transmission mechanisms:

- **Broadcast transmission**. An infected node spread the contagion across all its susceptible neighbors. The probability that *i* transmits to susceptible neighbor *k* under broadcast mode is given by:

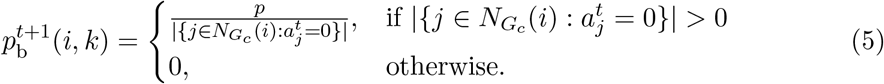
- **Targeted transmission**. An infected node preferentially targets the most context-vulnerable susceptible neighbor, such that ties are broken uniformly at random:

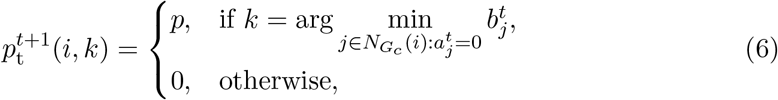

where 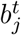 is the context score of susceptible individual *j*. Thus, lower context scores correspond to greater vulnerability to the contagion.

We set the transmission mode for the system at the beginning of the simulation. Let *m*_*S*_ ∈ {0, 1} determine whether the system uses broadcast (*m*_*S*_ = 0) or targeted (*m*_*S*_ = 1) transmission. Then, we define the probability of transmission of contagion by individual *i* to neighbor *k* at time *t* + 1 as:

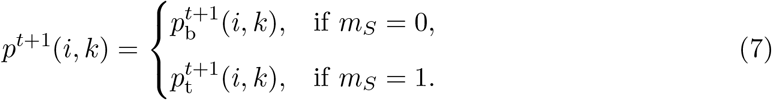

The local functions ℱ are probabilistic functions that determines the state of individual *i* at time *t*. We model the probability of infection of a susceptible individual by coupling the contagion spread with social bias of the individuals, namely:

- *Contagion Pressure:* We compute the contagion spread probability by neighbors of *i* Each infected neighbor attempts to transmit the individual independently, and the model computes the probability that at least one succeeds. This is expressed through a product term that represents the probability that all neighbors fail to infect *i*, and subtracting this quantity from one yields the net contagion spread probability i.e., contagion pressure.
- *Social Conditioning:* We compute the vulnerability of a susceptible individual to the contagions as a function of their social-context. A negative context renders an individual more vulnerable to infection, while positive makes them more resilient, with this modulation mapped smoothly into the interval [0, 1]. At 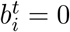, the conditioning becomes a fair coin toss.

We are now ready to define the effective probability of a susceptible node being infected at time *t*. Formally, if individual *i* is susceptible at time *t*, then:

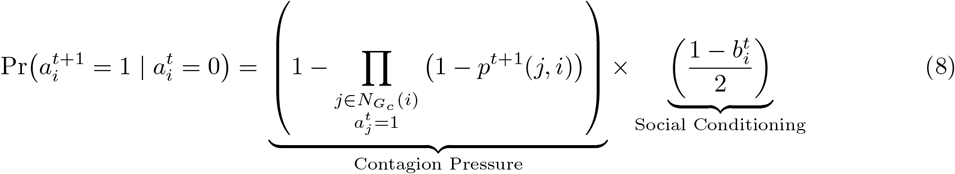

Figure 6 illustrates the state transitions for the individuals in the contagion layer, where the transition probabilities can be interpreted using Eq. (8). Note that state transitions occur discretely in the contagion state among {0, 1}, which is unlike the continuous updates in the contexts of the individuals.

**Figure 6:**
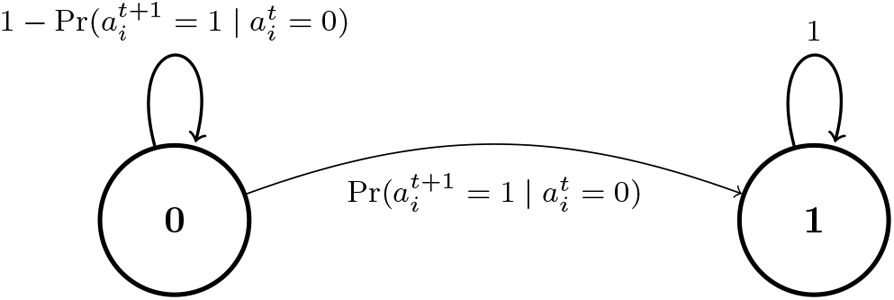
State-transition diagram for the contagion transmission layer of an individual *i*.

**Figure 7:**
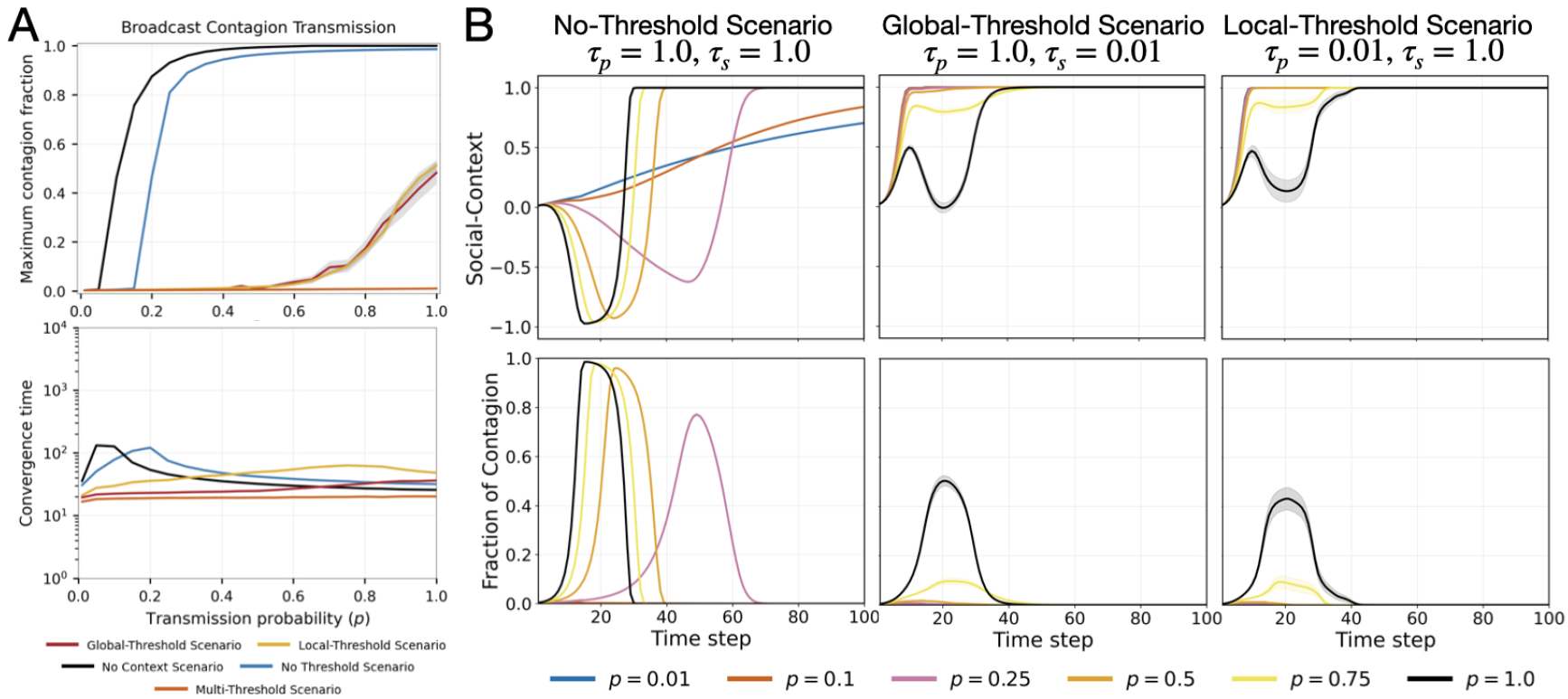
SIR contagion dynamics under broadcast transmission across context-thresholding scenarios. (A) We compare maximum contagion fractions and convergence time for the baseline SIR model, denoted by *No Context Scenario*, and four context-threshold variants. (B) We illustrate the evolution of the social-context layer (top row) and contagion prevalence (bottom row) for various transmission probabilities under the No-Threshold, Global-Threshold, and Local-Threshold scenarios.

### D Experimental Setup & Findings

In this section, we systematically discuss the framework parameter and their impact on the emergent contagion. All observations were averaged over 100 simulation runs with random seeds determining the placement of population characteristics.

#### System Setup

Our network consists of *n* = 30, 000 individuals and initial fraction of infected individuals to be 0.003. The individuals are part of two interaction graphs corresponding with the social-context layer and the contagion layer. The network is generated using the Barabási–Albert (BA) model with average degree 30 for the contagion layer. We augment the context layer network with 0.4% randomly inserted edges to the contagion layer topology.

**Table 1:** Framework Parameters.

| Parameter | Description | Baseline |
| --- | --- | --- |
| $n$ | Number of individual nodes in the network. | 30,000 |
| $G_S$ | Network Graph for Social-Context Layer of the system. | - |
| $G_C$ | Network Graph for Contagion Layer of the system. | - |
| $m_S$ | Transmission mode of the system. | - |
| $p$ | Transmission probability of contagion. | - |
| $f^+$ | Fraction of pro-social population at day 1. | 0.01 |
| $f^-$ | Fraction of infected population at day 1. | 0.003 |
| $\tau_p^l$ | Local-sensitivity threshold lower-limit for pro-social polarization. | $\tau_p$ |
| $\tau_p^u$ | Local-sensitivity threshold upper-limit for pro-social polarization. | 1 |
| $\tau_s^l$ | Global-Conformity threshold lower-limit for pro-social polarization. | $\tau_s$ |
| $\tau_s^u$ | Global-Conformity threshold upper-limit for pro-social polarization. | 1 |

### E Extension to SIR extension under broadcast transmission

We study SIR contagion dynamics under the broadcast transmission model within our framework. We consider two cases of recovery duration: 15 days and 180 days, after which they acquire immunity and cannot become susceptible again. To keep the modeling choice consistent across our framework, we treat recovered individuals as pro-social in the context layer; specifically, once an individual recovers, their context score is set to +1.

Under the SIR broadcast transmission model, social-context dynamics fundamentally reshape both the magnitude and temporal evolution of contagion spread. In the No-Threshold Scenario, contagion spreads rapidly and reaches high prevalence for sufficiently large transmission probabilities, while the social context gradually polarizes over time. In contrast, the Global-Threshold Scenario strongly suppresses contagion propagation, preventing large-scale outbreaks even at moderate transmission rates through coordinated shifts in collective social context. The Local-Threshold Scenario also reduces contagion prevalence, though suppression emerges more gradually through localized peer-driven adaptation. Across all regimes, recovered individuals transition into a permanently pro-social state, reinforcing social resistance after recovery and contributing to the long-term stabilization of the system.

Figure 8 illustrates SIR contagion dynamics with a 180-day recovery period. The extended recovery duration significantly increases both contagion prevalence and persistence, particularly in the No-Threshold Scenario, where outbreaks rapidly saturate the population even at low transmission probabilities. In contrast, the Global-Threshold and Local-Threshold scenarios suppress contagion spread through adaptive changes in the social-context layer. Panel B highlights the co-evolution of social context and contagion prevalence over time. The Global-Threshold regime exhibits strong nonlinear feedback and delayed transitions, while the Local-Threshold regime produces more gradual suppression through localized peer-driven resistance. Overall, the results demonstrate that prolonged infectious periods amplify the influence of adaptive social-context dynamics on outbreak evolution and stabilization.

**Figure 8:**
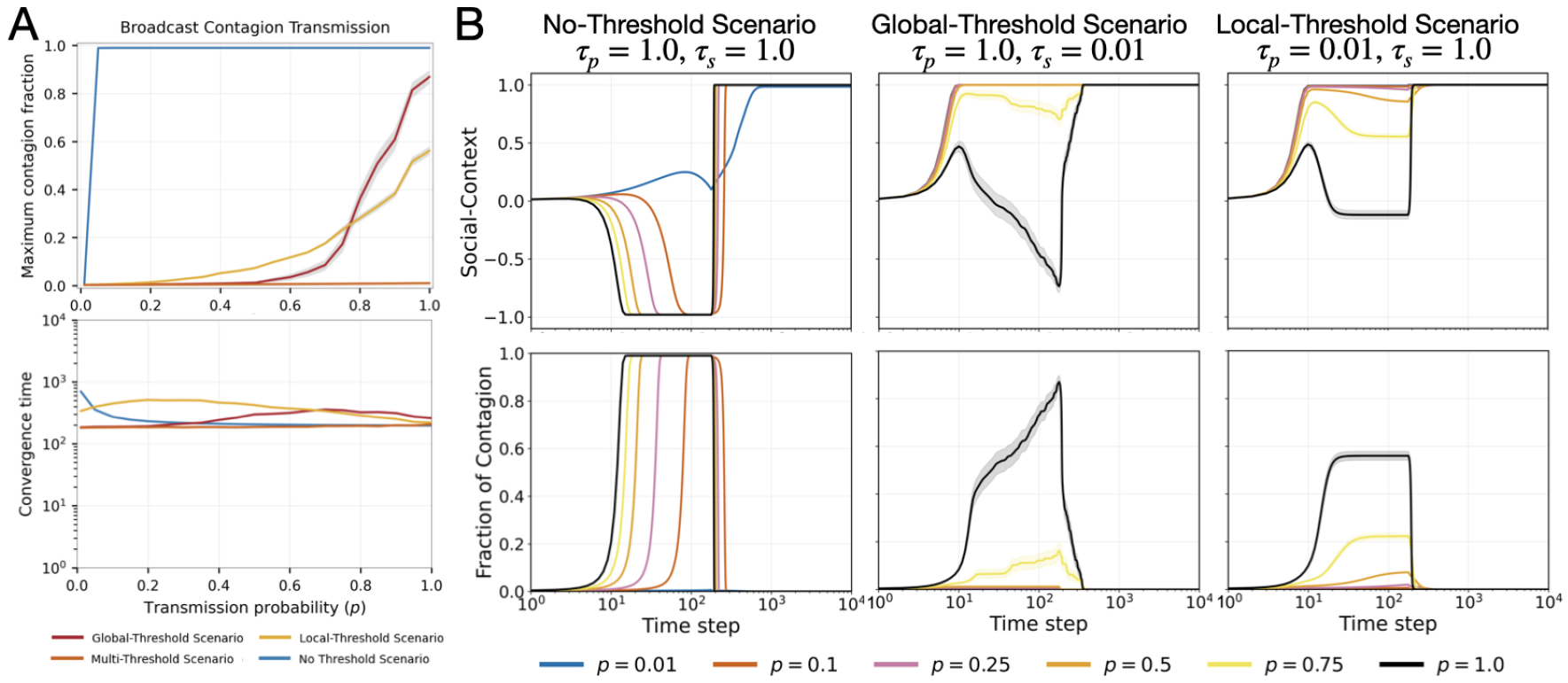
SIR contagion dynamics with 180 days recovery under broadcast transmission across context-thresholding scenarios. Note the log scale of the x-axis in (B).

**Figure 9:**
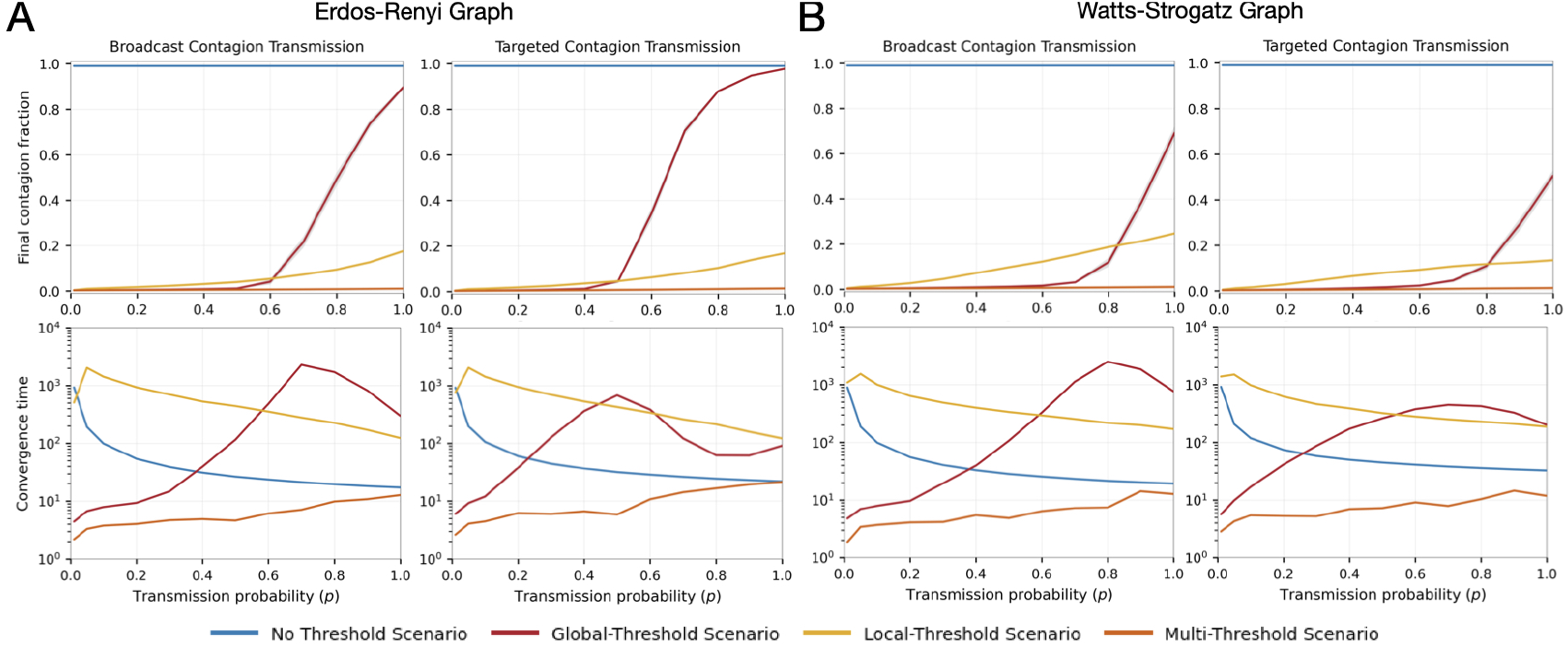
Impact of network topology on the contagion dynamics. We compare the final contagion fraction and convergence time for the Erdos-Renyi Graph and Watts-Strogatz Graph at average node degree of 30.

Compared to the 15-day recovery setting, the 180-day recovery regime produces substantially larger and longer-lasting outbreaks. With shorter recovery times, infected individuals quickly transition into recovered pro-social states, allowing social resistance to accumulate rapidly and suppress contagion spread. In contrast, the longer recovery delays this stabilizing feedback, enabling contagion to persist and propagate for extended periods before recovery-driven social reinforcement emerges. This effect is especially pronounced in the No-Threshold and Global-Threshold scenarios, where outbreaks exhibit sustained high prevalence and significantly slower convergence. These results highlight how recovery duration critically shapes the balance between contagion amplification and adaptive social resistance.

### F Additional Empirical Results

In this section, we present supporting results that complements the findings int he main paper. We study impact of varying network topology with comparable individual connectivity property i.e., average node degree in Section F.1. Then, we illustrated the temporal dynamics of multi-threshold scenario under high heterogeneity in Section F.2.

#### F.1 Contagion dynamics under comparable network topology

At a broad level, we observe that the emergent contagion dynamics under alternative network topologies closely mirror those observed in Fig. 2. Despite varying the underlying graph generation model while maintaining comparable average node degree, the coupled framework consistently reproduces the same qualitative dynamical regimes, including suppression, delayed activation, and sharp nonlinear transitions across the different thresholding scenarios.

In particular, the Erdős-Rényi topology exhibits contagion dynamics most similar to the baseline behavior observed in Fig. 2, especially under targeted contagion transmission. The final contagion fraction and convergence profiles under the Global-Threshold and Multi-Threshold Scenarios closely replicate the phase-transition behavior observed in the original network setting, suggesting that the emergent tipping dynamics are robust to random graph connectivity structure when neighborhood size remains comparable.

In contrast, the Watts–Strogatz topology produces more pronounced deviations under broadcast contagion transmission, especially in the Local-Threshold Scenario. Relative to Fig. 2, contagion affects a larger fraction of the network, and convergence times remain elevated across a broader range of transmission probabilities. This behavior is likely driven by the short path lengths characteristic of Watts–Strogatz networks, which allow broadcast transmission to expose susceptible individuals through many rapidly reachable pathways. At the same time, because the Local-Threshold Scenario activates resistance based on each individual’s local social context, rapid diffusion over a low-diameter network exposes individuals to a more heterogeneous set of neighboring states. This can keep the social context in an intermediate regime for longer, delaying threshold activation and prolonging exposure to contagion before the system converges. Notably, even under the Global-Threshold Scenario, where final contagion prevalence remains low relative to the other two network families at comparable average degree, convergence times remain elevated at higher transmission probabilities. This suggests that heterogeneous exposure can delay stabilization even when global social feedback ultimately suppresses widespread contagion.

More broadly, these results suggest that the qualitative contagion regimes induced by heterogeneous social-context depend more strongly on the scale and structure of social feedback mechanisms than on the specific graph generation model itself. While network topology modulates the sharpness of transitions and the efficiency of spreading pathways, the coupled framework consistently preserves the core emergent phenomena across graph families when local neighborhood size remains comparable.

#### F.2 Temporal dynamics of Multi-Threshold Scenario under High Heterogeneity

Under highly heterogeneous thresholding with *τ*_*p*_ = *τ*_*s*_ = 0.01, contagion remains strongly suppressed and converges rapidly to a low-prevalence equilibrium. As illustrated in Fig. 10, the contagion fraction increases only marginally before stabilizing within a few time steps, while the social-context rapidly aligns toward a strongly pro-social state due to context diffusion. The substantial heterogeneity in local and global thresholds generates persistent social barriers that prevent large-scale synchronization of contagion spread, effectively containing transmission before widespread cascades can emerge. Consequently, the system quickly settles into a stable suppression regime characterized by low contagion prevalence and rapid convergence dynamics.

**Figure 10:**
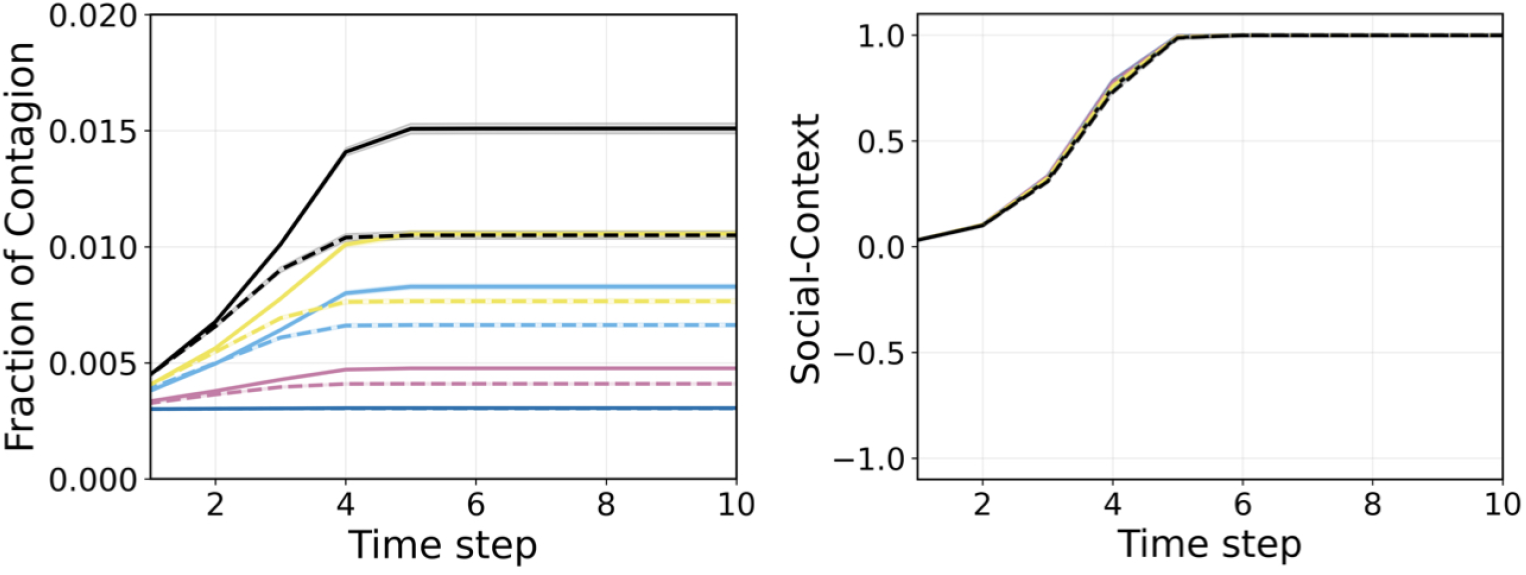
Evolution of Contagion and Social-Context under highly heterogenous multi-threshold scenario with *τ*_*p*_ = *τ*_*s*_ = 0.01.

**Figure 11:**
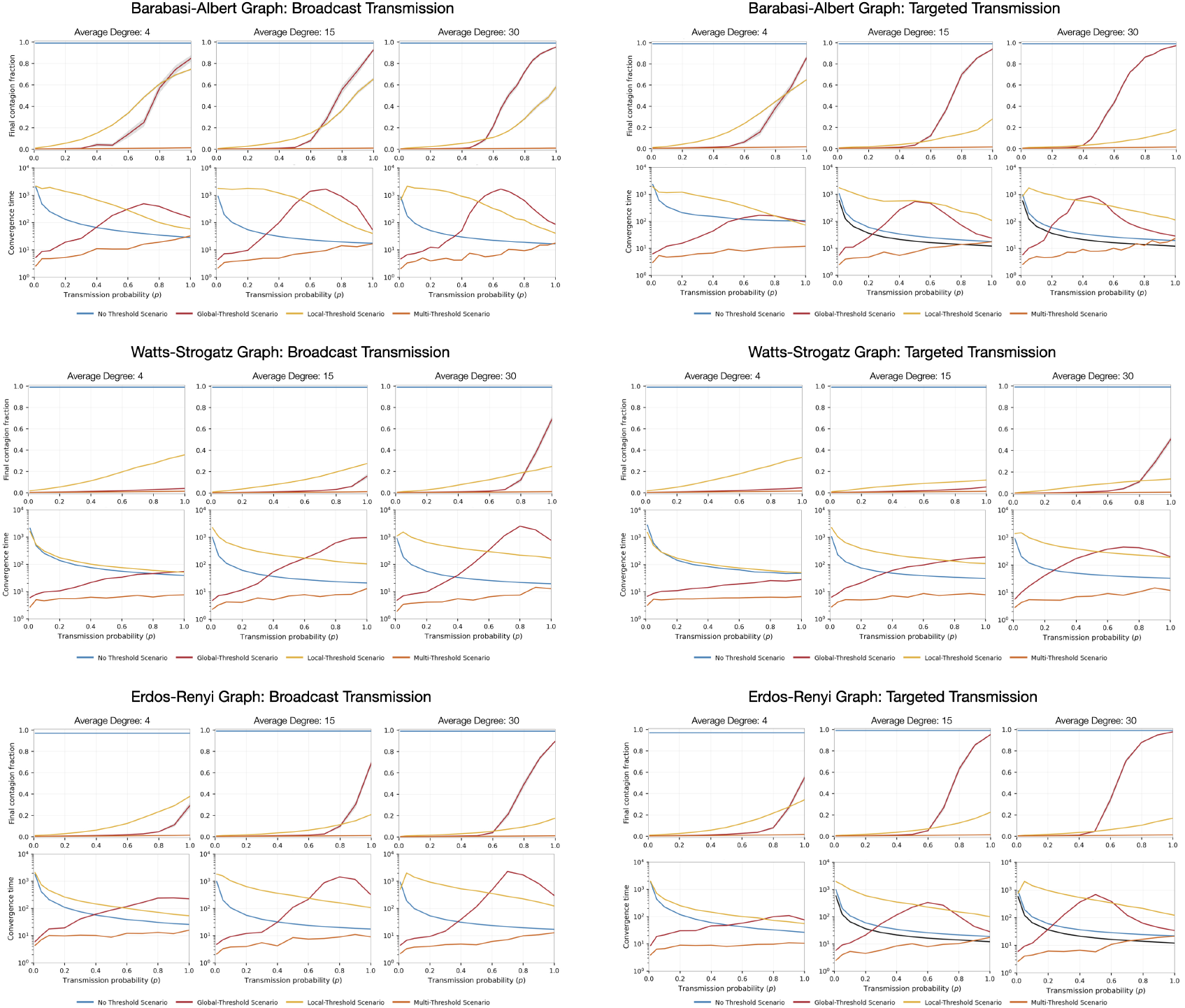
Robustness of co-evolving contagion dynamics across network topologies and connectivity regimes.

#### F.3 Robustness to Network Topology

We evaluate the impact of network structure on the emergent contagion dynamics under the proposed context-coupled framework across three graph families: Barabási-Albert, Watts-Strogatz, and Erdős-Rényi networks. For each topology, we compare broadcast (left) and targeted (right) contagion transmission mechanisms under varying average network degree. The top row within each panel illustrates the final contagion fraction at equilibrium as a function of transmission probability *p*, while the bottom row reports the corresponding convergence time.

Across all network families, the coupled framework consistently produces suppression regimes, delayed activation, and nonlinear contagion transitions induced by heterogeneous social-thresholding. Increasing network connectivity generally sharpens phase transitions and accelerates contagion spread once activation thresholds are crossed, particularly under the Global-Threshold Scenario. In contrast, the Local-Threshold and Multi-Threshold Scenarios maintain partial contagion suppression and elevated convergence times across scenarios, reflecting persistent resistance generated by social feedback.

## References

[1] Ali, M. M., Amialchuk, A., and Dwyer, D. S. The social contagion effect of marijuana use among adolescents. PloS one 6, 1 (2011), e16183.

[2] Barbieri, N., Bonchi, F., and Manco, G. Topic-aware social influence propagation models. Knowledge and information systems 37,3 (2013), 555–584.

[3] Barsade, S. G. The ripple effect: Emotional contagion and its influence on group behavior. Administrative science quarterly 47, 4 (2002), 644–675.

[4] Calvo, S., Carrasco, J. P., Conde-Pumpido, C., Esteve, J., and Aguilar, E. J. Does suicide contagion (werther effect) take place in response to social media? a systematic review. Spanish journal of psychiatry and mental health (2024).

[5] Campbell, E., and Salathé, M. Complex social contagion makes networks more vulnerable to disease outbreaks. Scientific Reports 3 (2013), 1905.

[6] Cencetti, G., and Battiston, S. Diffusive behavior of multiplex networks. New Journal of Physics 21, 3 (2019), 035006.

[7] Centola, D., and Macy, M. Complex contagions and the weakness of long ties. American Journal of Sociology 113, 3 (2007), 702–734.

[8] Dodds, P. S., and Watts, D. J. Universal behavior in a generalized model of contagion. Physical Review Letters 92, 21 (2004), 218701.

[9] Dodds, P. S., and Watts, D. J. A generalized model of social and biological contagion. Journal of Theoretical Biology 232, 4 (2005), 587–604.

[10] Epstein, J. M., Parker, J., Cummings, D., and Hammond, R. A. Coupled contagion dynamics of fear and disease. PLoS ONE 3, 12 (2008), e3955.

[11] Fu, F., Rosenbloom, D. I., Wang, L., and Nowak, M. A. Imitation dynamics of vaccination behavior on social networks. Proceedings of the Royal Society B 278, 1702 (2011), 42–49.

[12] Funk, S., Gilad, E., Watkins, C., and Jansen, V. A. A. The spread of awareness and its impact on epidemic outbreaks. Proceedings of the National Academy of Sciences USA 106, 16 (2009), 6872–6877.

[13] Granell, C., Gómez, S., and Arenas, A. Dynamical interplay between awareness and epidemic spreading. Physical Review Letters 111, 12 (2013), 128701.

[14] Granovetter, M. Threshold models of collective behavior. American journal of sociology 83, 6 (1978), 1420–1443.

[15] Gross, T., D’Lima, C. J. D., and Blasius, B. Epidemic dynamics on an adaptive network. Physical Review Letters 96, 20 (2006), 208701.

[16] Herrera-Diestra, J. L., and Meyers, L. A. Local risk perception enhances epidemic control. PloS one 14, 12 (2019), e0225576.

[17] Howard, J., Huang, A., Li, Z., Tufekci, Z., Zdimal, V., Van Der Westhuizen, H.-M., Von Delft, A., Price, A., Fridman, L., Tang, L.-H., et al. An evidence review of face masks against covid-19. Proceedings of the National Academy of Sciences 118, 4 (2021), e2014564118.

[18] Huang, C.-W., Hu, T., Zheng, H., Wu, Y.-L., Li, J.-M., Wang, Y.-M., Su, W.-J., Wang, W., Liu, Y.-Z., and Jiang, C.-L. Contagion of depression: a double-edged sword. Translational Psychiatry 14, 1 (2024), 396.

[19] Huang, W., Zhang, X., Zhu, X., and Chen, G. Role of persuasion in social contagion. Physica A 457 (2016), 179–189.

[20] Iyengar, R., Van den Bulte, C., and Valente, T. W. Opinion leadership and social contagion in new product diffusion. Marketing science 30, 2 (2011), 195–212.

[21] Jovanovski, P., Kocarev, L., and Stadler, P. F. Multilayer SIS model with multiple contagions. Physical Review E 98, 2 (2018), 022310.

[22] Kaplan, E. H. A method for evaluating needle exchange programmes. Statistics in medicine 13, 19-20 (1994), 2179–2187.

[23] Kempe, D., Kleinberg, J., and Tardos, É. Maximizing the spread of influence through a social network. In Proceedings of the ninth ACM SIGKDD international conference on Knowledge discovery and data mining (2003), pp. 137–146.

[24] Krassa, M. A. Social groups, selective perception, and behavioral contagion in public opinion. Social Networks 10, 2 (1988), 109–136.

[25] Kremer, L., Holkenborg, S. K., Reimert, I., Bolhuis, J., and Webb, L. The nuts and bolts of animal emotion. Neuroscience & Biobehavioral Reviews 113 (2020), 273–286.

[26] Kuhlman, C. J., Tuli, G., Swarup, S., Marathe, M. V., and Ravi, S. S. Blocking simple and complex contagion by edge removal. In Proceedings of the IEEE/ACM International Conference on Advances in Social Networks Analysis and Mining (2015).

[27] Lee, J., Yu, F., Auyong, H., and Chok, S. Community-based approaches to the prevention, rehabilitation and reintegration of drug offenders, 2018.

[28] Liang, Z., He, Q., Du, H., and Xu, W. Targeted influence maximization in competitive social networks. Information Sciences 619 (2023), 390–405.

[29] Myers, S. A., Zhu, C., and Leskovec, J. Information diffusion and external influence in networks. In Proceedings of the 18th ACM SIGKDD international conference on Knowledge discovery and data mining (2012), pp. 33–41.

[30] Newman, M. E. Spread of epidemic disease on networks. Physical review E 66, 1 (2002), 016128.

[31] Qian, M., and Jiang, J. Covid-19 and social distancing. Journal of Public Health 30, 1 (2022), 259–261.

[32] Qiu, Z., Espinoza, B., Vasconcelos, V. V., Chen, C., Constantino, S. M., Crabtree, S. A., Yang, L., Vullikanti, A., Chen, J., Weibull, J., Basu, K., Dixit, A., Levin, S. A., and Marathe, M. V. Understanding the co-evolution of mask-wearing and epidemics: A network perspective. Proceedings of the National Academy of Sciences USA 119, 5 (2022), e2123355119.

[33] Ramazi, P., Riehl, J., and Cao, M. Networks of conforming and anti-conforming agents. Games and Economic Behavior 96 (2016), 86–102.

[34] Salathé, M., and Bonhoeffer, S. The effect of opinion clustering on disease outbreaks. Journal of the Royal Society Interface 7, 55 (2010), 1505–1508.

[35] Sampson, C. R., and Restrepo, J. G. Competing social contagions with opinion-dependent infectivity. Physical Review E 111, 2 (2025), 024313.

[36] Thu, T. P. B., Ngoc, P. N. H., Hai, N. M., and Tuan, L. A. Effect of the social distancing measures on the spread of covid-19 in 10 highly infected countries. Science of the Total Environment 742 (2020), 140430.

[37] van den Ende, M. W., van der Maas, H. L., Epskamp, S., and Lees, M. H. Alcohol consumption as a socially contagious phenomenon in the framingham heart study social network. Scientific Reports 14, 1 (2024), 4499.

[38] Watts, D. J. A simple model of global cascades on random networks. Proceedings of the National Academy of Sciences USA 99, 9 (2002), 5766–5771.

[39] Zhang, R., Wang, X., and Pei, S. Targeted influence maximization in complex networks. Physica D: Nonlinear Phenomena 446 (2023), 133677.

